# Individual variation of natural *D. melanogaster* associated bacterial communities

**DOI:** 10.1101/126912

**Authors:** Yun Wang, Fabian Staubach

## Abstract

*D. melanogaster* has become an important model organism to study host-microbe interaction. However, we still know little about the natural microbial communities that are associated with *D. melanogaster.* Especially, information on inter-individual variation is still lacking because most studies so far have used pooled material from several flies. Here, we collected bacterial 16S rRNA gene community profiles from a set of 32 individuals from a single population and compare the variation to that of samples collected from different substrates and locations. While community differences were on average larger between samples collected from different substrates, there was still a surprising amount of variation of microbial communities between individual flies. The samples clustered into two groups suggesting that there are yet unknown factors that affect the composition of natural fly associated microbial communities and need research.

**Importance:** *D. melanogaster* is an important model organism in evolutionary biology and also for the study of host-microbe interaction. In order to connect these to aspects of *D. melanogaster* biology, it is crucial to better understand the natural *D. melanogaster* microbiota because only the natural microbiota can affect the evolution of the host. We present, to our knowledge, the first data set that captures inter-individual variation of *D. melanogaster* associated bacterial communities. Clustering of communities into two larger groups suggests that there are important drivers of these communities that we do not understand yet suggesting in return that more research on the natural microbiota of *D. melanogaster* is needed.

## Introduction

*Drosophila melanogaster* has become an important model for the investigation of host-microbe interaction (1). Interactions with bacteria can affect *D. melanogaster* phenotype in many aspects. Bacteria mediated *D. melanogaster* phenotypic effects range from pathogenic (2), over effects on cold tolerance (3), and effects on *D. melanogaster* nutritional status (4, 5) to highly beneficial effects. For example, certain acetic acid bacteria (Acetobacteraceae) and Lactobacillaceae significantly promote *D. melanogaster* larval growth on amino acid poor diet (6, 7) and can ensure longtime fertility and longevity under nutrient poor conditions (8). Because bacteria from the same families that have these highly fitness relevant effects on *D. melanogaster* can also be found in association with wild-caught flies (9–12), it seems reasonable to assume that they could also play a role in fly evolution. However, in most studies bacterial strains and communities that were isolated from lab-reared flies are investigated and we still know rather little about natural *Drosophila* associated bacterial communities and the factors shaping them. Yet, only natural microbial communities can have played a role in *Drosophila* evolutionary history. This is the more important since it was shown by Chandler et al. (2011) (11) and Staubach et al. (2013) (12) that bacterial communities associated with wild-caught flies are different from those associated with lab-reared flies. Under controlled laboratory conditions, host genetic make up can influence *D. melanogaster* associated bacterial communities (5, 13). Under natural conditions, the substrate flies were collected from strongly correlates with bacterial community composition (12), while species differences between *D. melanogaster* and *D. simulans* are detectable but have a smaller effect.

These studies on natural microbial communities relied on pooling material from several flies for bacterial community profiling. However, information from individual, wild-caught flies is required to put the size of substrate related effects on communities into the perspective of microbial community variation between individuals. A better understanding of inter-individual variation within populations can also help us to assess if the current practice to pool material from several individuals is appropriate to represent the microbial community of a local population. Finally, bacterial community profiles of individuals collected from the same diet at the same time could help to evaluate whether there are other factors beyond diet that drive variation in natural fly associated microbial communities. Yet, contrary to humans, where collecting individual microbial profiles has been in the focus (14), such data, to our knowledge, does not exist for *D. melanogaster.*

In order to assess bacterial community variability between individuals under otherwise constant conditions, we collected flies from the same substrate and location at the same time and assessed the bacterial communities of individual male flies using 16S rRNA gene profiling. We compared and contrasted these communities to communities from flies collected from different populations and substrates to place variation between individuals into the context of variation caused by known factors that influence microbial communities. Furthermore, we assess and evaluate how the common practice of pooling material from several individuals influences diversity measurements and composition of fly associated microbes.

## Results

We assessed bacterial community composition and diversity by sequencing the V4 region of the 16S rRNA gene for 32 individual and a pool of 5 wild-caught D. *melanogaster* that were collected from the same substrate (plums) at the same location and time. Additionally a pool of 5 male flies from the same location and substrate, but collected 1 year earlier was analyzed. In order to assess between population variation 11 pools of flies collected from 7 different substrates and locations were analyzed. All pools were based on 5 flies except Orange3, for which we were able to obtain only 3 flies. Orange3 was no outlier in any of the analyses and was included with the pools of 5 flies (see Table S1 for an overview of all samples). A total of ∼ 2,240,000 sequences passed quality filtering. ∼1,050,000 sequences matched the *Wolbachia* 16s rRNA gene sequence and were removed. Individual fly sample #6 was removed because only 11 sequences remained after *Wolbachia* removal. At least 1008 sequences per sample were collected for the remaining samples.

### Bacterial diversity of pools and individual flies is similar

We were interested in finding out whether individual flies carry reduced bacterial diversity or a skew in bacterial abundance patterns, as might be expected from stronger stochastic effects in smaller samples, when compared to pools of 5 flies. For comparing bacterial alpha diversity between individual flies and pools of 5 flies, we grouped sequences into 97% identity OTUs (Operational Taxonomic Units).

The mean number of OTUs observed when sampling 1008 sequences did not differ between individual flies (44.4±17.8 OTUs) and the pool of 5 flies (40 OTUs) that was collected from the same substrate at the same time (*P* = *1*, Mann-Whitney test, Figure 1A, Table 1). This indicates that bacterial OTU richness of the plum population is well represented by a pool of 5 flies. Surprisingly, OTU richness of pools of 5 flies collected across substrates and locations also did not differ from the average richness found associated with individual flies collected from plums (39.3±23.8 OTUs, *P* = 0.23, Mann-Whitney test), providing no evidence for a sample size related reduction in diversity. The same holds true for Chao’s richness estimate (15), Shannon’s H (16), and Simpsons D (17) (Table 1). Furthermore, there was also no difference in variance of OTU richness between communities from individual flies and that of pools of 5 (Levene-test, *P* = 0.21, Figure 1B) indicating that the variance in bacterial richness between individuals from the same population and substrate can be similar to that between *D. melanogaster* populations from different substrates and locations.

**Figure 1.**
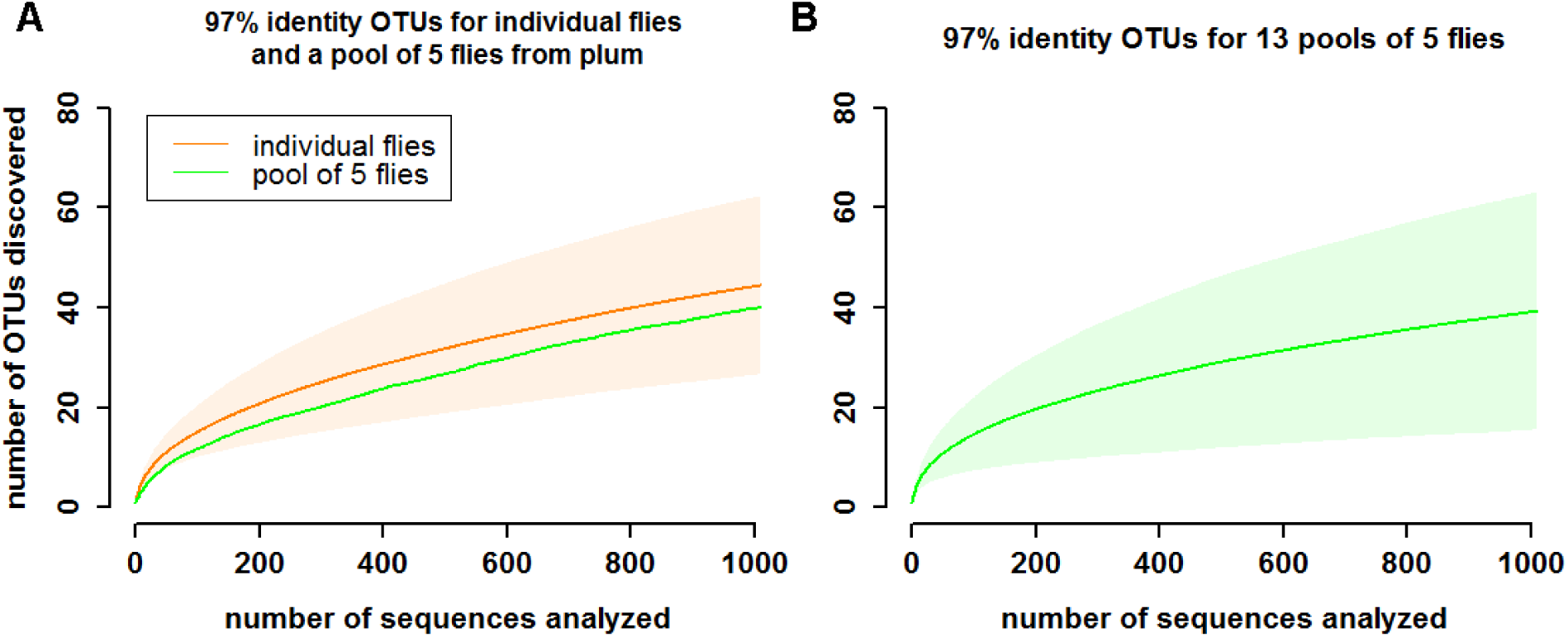
Rarefaction curves of 97% identity OTUs (A) for individual flies and a pool of 5 flies from the same substrate (plum). The orange line shows the mean number of OTUs discovered for 31 individual fly samples with the shaded area indicating the standard deviation. The green line represents a pool of 5 flies sampled from the same substrate. (B) for 13 pools of five flies collected across substrates and locations. The green line represents the mean and the shaded area the standard deviation.

**Table 1.** Alpha diversity of individual and pooled samples

| sample | #OTU | P | Chao | P | Shannon H | P | Simpson D | P | Shannon E | P | Simpson E | P |
| --- | --- | --- | --- | --- | --- | --- | --- | --- | --- | --- | --- | --- |
| individual flies | 44.4±17.8 |  | 75.9±32.0 |  | 1.94±0.62 |  | 0.29±0.20 |  | 0.51±0.13 |  | 0.11±0.04 |  |
| pools of 5 from plum | 40 | 1 | 63.8 | 0.81 | 1.17 | 0.31 | 0.57 | 0.31 | 0.32 | 0.19 | 0.04 | 0.13 |
| pools of 5 | 39.3±23.7 | 0.23 | 71.2±55.6 | 0.27 | 1.89±0.81 | 0.93 | 0.31±0.25 | 0.93 | 0.52±0.18 | 0.57 | 0.13±0.04 | 0.11 |

Analysis of evenness using Shannon’s E and Simpson’s E also revealed no significant differences between individuals and pools of 5 flies. This indicates that there are no significant skews introduced into the OTU distribution by sampling individuals when compared to pools of 5 flies.

### Bacterial communities are dominated by acetic acid bacteria and vary in composition between individuals from the same population

For a detailed view of the variability in bacterial community composition of individual and pooled fly samples, we classified the 16S rRNA gene sequences taxonomically.

The bacterial communities are dominated by acetic acid bacteria (Acetobacteraceae 62.6%) representing the 4 of the 5 most common genera (*Saccharibacter* 24.1%, *Gluconobacter* 18.1%, *Acetobacter* 13.4%, and *Gluconacetobacter* 6.7% average relative abundance). The relative abundance of the different taxa is highly variable between individuals from the same population (*Saccharibacter* 2.0% - 93.9% relative abundance, *Gluconobacter* 1.7% - 64.9%, *Acetobacter* 0.5% - 37.2%, and *Gluconacetobacter* 0.4% - 43.9%). Please note that bootstrap support for the *Saccharibacter* classification is often relatively low (between 40% and 60%, Table S2) indicating that there are several sequences in the SILVA database that match sequences from this taxonomic group similarly well. Blast search for representative sequences from this taxonomic group produced perfect matches to bacteria classified as *Commensalibacter* in Chandler et al. (2011) (11) and *Acetobacter* in Corby-Harris et al. (2007) (9) (see File S1 for blast search results). Acetic acid bacteria represent a major bacterial group associated with wild-caught flies (9, 11, 12). Enterobacteriaceae are also common (16.8% average relative abundance) with *Buttiauxella* (8.2% average relative abundance) and *Serratia* (6.8% average relative abundance) being the most common. Members of the genus *Serratia* can be *Drosophila* pathogens and occur at high relative abundance in individual samples. This pattern of low abundance or absence in most samples and sharp increase in individual samples (sample orange2, 44.0% rel. abundance) has been associated with *Drosophila* pathogens (12). A similar pattern is visible for *Enterococcus* in sample orange1 (46.5% rel. abundance compared to 6.9% average relative abundance) and the tomato sample (53.6% rel. abundance). *Enterococcus* can reach high titers in *Drosophila* and cause mortality (10).

### Beta diversity between individual samples is smaller than between pools collected across locations and substrates

As described above, microbial community composition varies substantially even between individual flies from the same population. In order to relate this variation between individuals to variation between populations, we analyzed beta-diversity using Bray-Curtis community distances (BCD) and included samples from different substrates and locations in the analysis.

The pairwise BCD between samples from individual flies is smaller than that of pools of 5 flies sampled across substrates and locations (Figure 3, *P* = 2.3 x 10^-10^, Mann-Whitney-test) indicating that between population variation is larger than within population variation. However, the difference is small (0.15) and the BCDs between populations fall almost completely within the range of distances between individuals. These results hold true for Jaccard (Figure S1) as well as weighted (Figure S2) and unweighted (Figure S3) Unifrac community distance.

**Figure 3.**
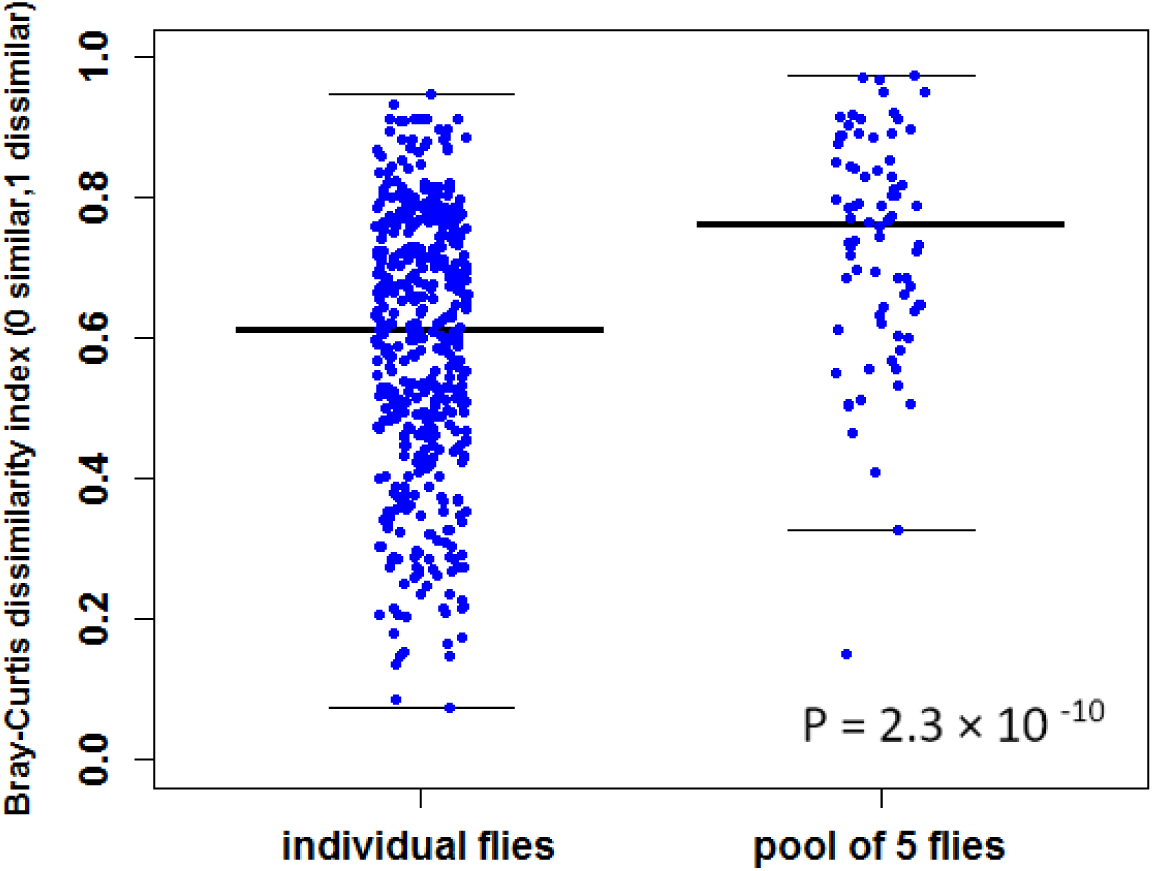
Pairwise Bray-Curtis distance for individual and pooled samples. P-value was computed using the Mann-Whitney-U-test.

### Individual fly samples from the same population fall into two groups

In order to explore potential factors shaping fly associated bacterial communities, a Principal Coordinate Analysis was carried out using BCD (Figure 4).

**Figure 4.**
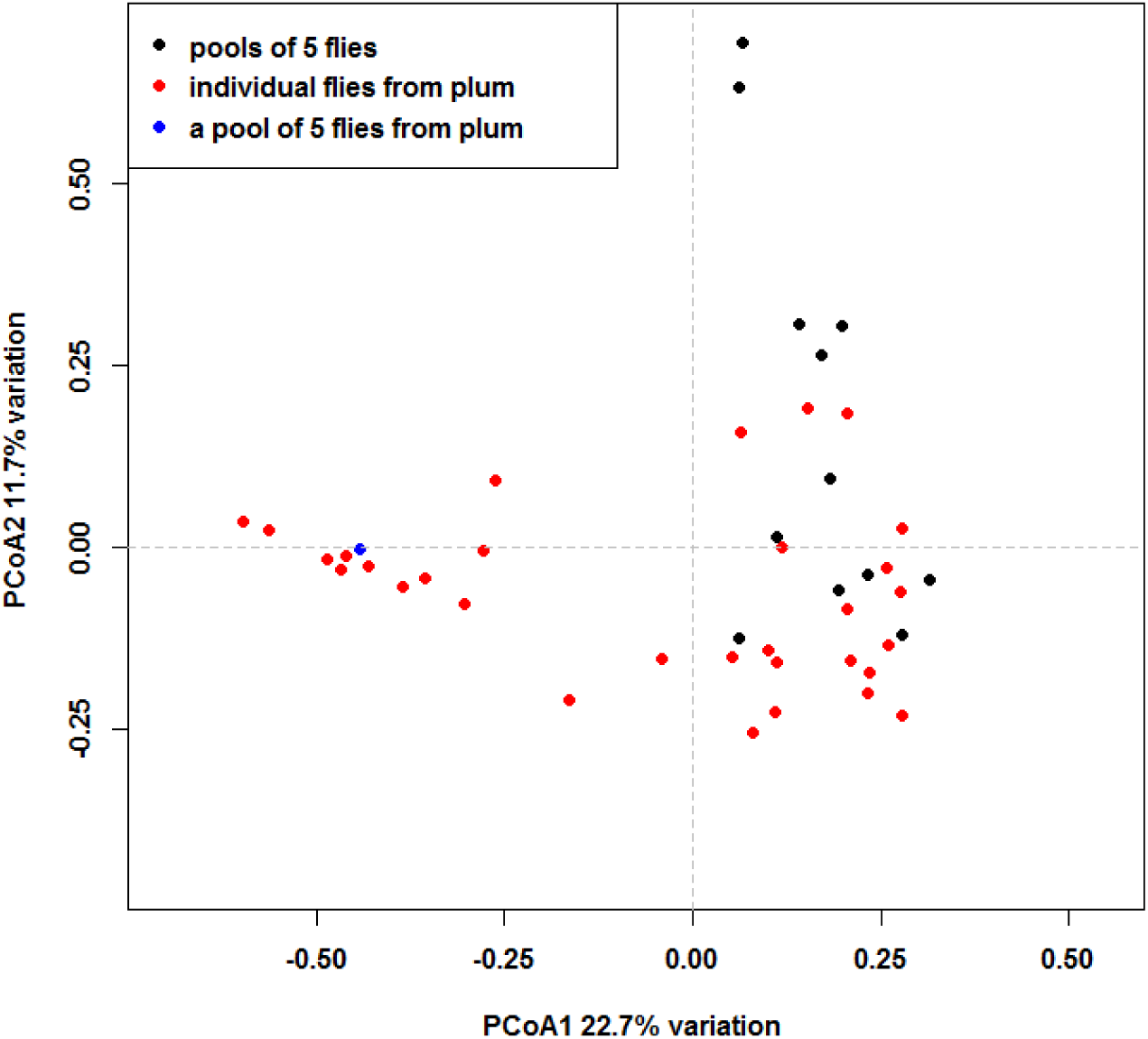
Principal Coordinates Analysis of Bray-Curtis distances.

In this analysis most of the variation between individual fly samples falls onto the first PCo, while the pools of 5 flies are distributed along the second PCo2. The two samples collected from grapes that have a high relative abundance of *Buttiauxella* are apart from the other samples at the top of the graph. Interestingly, Figure 4 suggests that the individual fly samples fall into two groups. The pool of 5 flies from plums is at the center of one of these groups suggesting that it represents only a part of the variation found in the plum population.

These two groups are supported by hierarchical clustering based on BCD (Figure 5), Jaccard-Distances (Figure S4), and weighted Unifrac distances (Figure S5), but not by unweighted Unifrac distances (Figure S6). In order to identify signature taxa that contribute to differences between the two groups, we applied the SIMPER method combining the samples in the two groups into two different habitats. This identifies OTU1 that was classified as *Saccharibacter* as the largest contributor (19.8% of variation) to the dissimilarity between two groups (a list of contributing OTUs can be found in Table S3).

**Figure 5.**
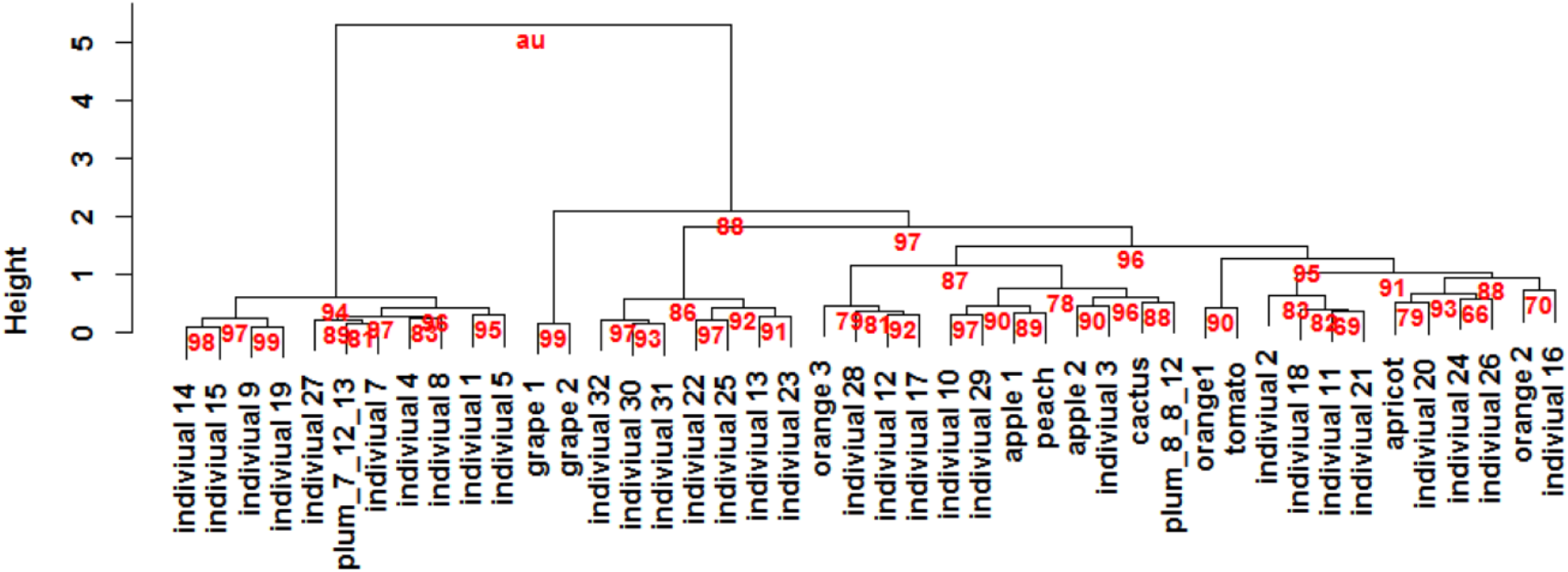
Hierarchical clustering of all samples based on Bray-Curtis dissimilarities. Values at branches are AU (Approximately Unbiased) bootstrap support (31).

## Discussion

In this study we described bacterial community variation between individual wild-caught *D. melanogaster.* By analyzing a sample of 32 individual flies from the same population, substrate, and at the same time, we were able to address the questions (i) whether the common practice of pooling material from several flies for bacterial community analysis leads to representative community assessment and (ii) whether there is evidence for factors other than the substrate the flies were collected from that shape natural bacterial communities, and (iii) compare within population variation of bacterial communities to between population variation. As before (12), we used entire flies for our study. This is important because fly pathogens can reach high titers in the hemolymph (2, 10) and might be overlooked by focusing on the gut. Although bacteria on the fly surface contribute only ∼10% to the total bacterial load of flies (18), and hence their effect on bacterial community composition should be minor, they could still play a role in inoculating fruit and shaping the microbial environment of *D. melanogaster.*

### Alpha diversity of individual fly and pooled fly communities

Bacterial diversity varied extensively between individuals from the same substrate and location with the standard deviation of OTUs discovered at 40% of the mean. Pooling of 5 individuals from the same population allowed for a good estimate of the mean OTU richness and evenness of individual samples. Interestingly, the variation in richness and evenness between individuals from the same population was as large as between populations that were collected from different locations and substrates. This further indicates that variation in community diversity between individuals of a population can be rather large.

Shannon’s diversity (H = 1.94 +/- 0.62) was comparable to that found associated with wild-caught flies in Staubach et al. 2013 (12) (H = 1.79 +/- 0.44) and hence to that from Chandler et al. 2011 (11) and Cox and Gilmore (2007) (10) who also investigated bacterial communities of wild-caught flies. Please see Staubach et al. (2013) (12) for extensive diversity comparisons between studies.

### Composition of bacterial communities

Concordant with many other studies acetic acid bacteria dominated the bacterial communities (10–12, 19). Also concordant with earlier studies on wild-caught flies enterobacteria occur at sometimes high relative abundance in some samples. This pattern has been connected to pathogens before and could indicate systemic infections (12).

We were surprised to find sequences classified as *Halomonas* in our samples as this is a halophile neither expected on rotting fruit nor flies. Blast search of a representative sequence from the largest *Halomonas* OTU identified plant mitochondria as best hits (see File S2 for blast search). The occurrence of plant mitochondrial sequence in our samples appears much more likely than *Halomonas.*

A representative sequence of the unclassified gammaproteobacterium (grey in Figure 2) that can be found in several samples matches perfectly with sequences from uncultured enterobacteria isolated from social corbiculate bees and nectar feeding bats (file S3). The high sugar content that nectar and rotting fruit have in common might favor the growth of similar bacteria.

**Figure 2.**
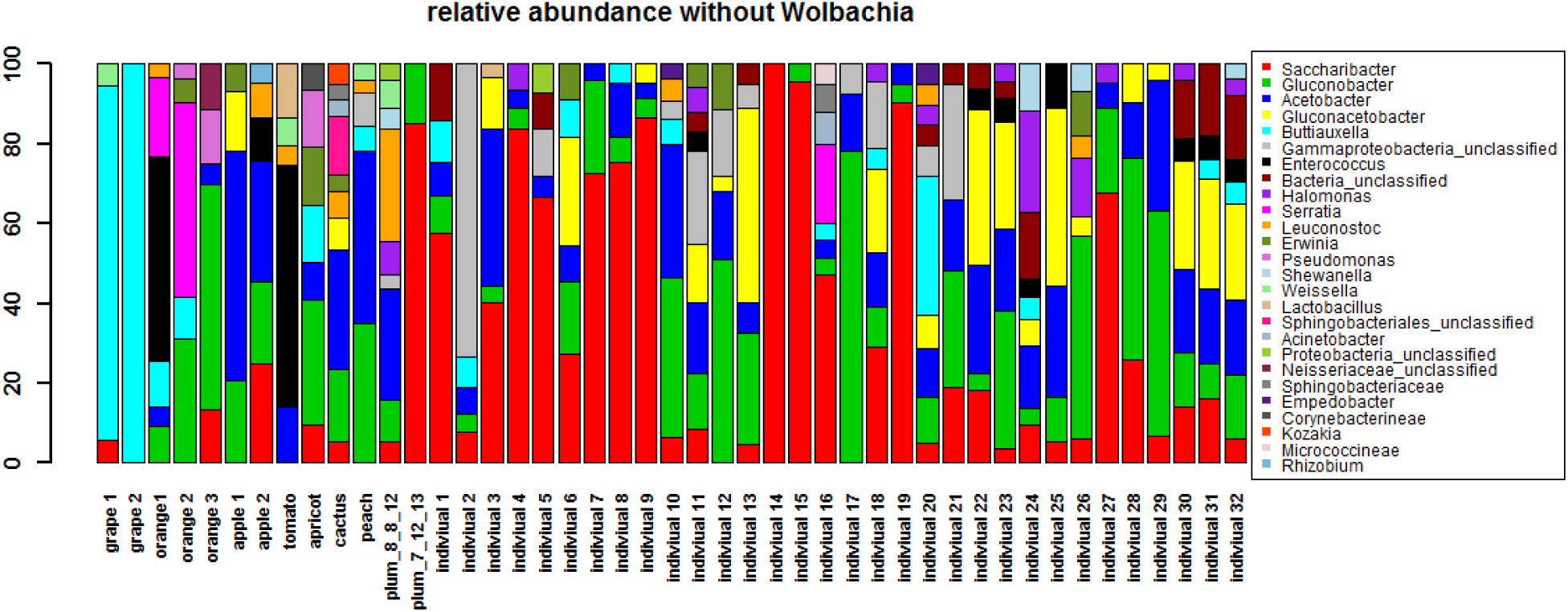
Relative abundance of bacterial taxa as assessed by 16s rRNA gene sequences. *Wolbachia* sequences were excluded. Bacterial genera of abundance <3% have been removed for clarity.

A full list of the 25 most common OTUs and representative sequences can be found in Table S2.

### Beta diversity

Bacterial community distances are on average higher between samples that were collected across different substrates and locations than between individual flies from a single population that was sampled at one point in time. This is not surprising since it has been shown before that substrate or a correlating variable, for example season, is an important factor for bacterial community composition associated with *D. melanogaster* and *D. simulans* (12). Nonetheless, the community distances between individual flies can be high and the distributions of pairwise community distances largely overlap between individuals and populations revealing surprising variability within a population. Assuming that flies continuously exchange microbes with their environment and that they replenish their gut microbiota through uptake of environmental bacteria (20) the large variation of bacterial communities of individuals could result from heterogeneity of the plum substrate the flies were collected from.

Surprisingly, our samples clustered into two groups. The clustering correlated with the most abundant OTU (OTU1, supposedly *Saccharibacter*). We can only speculate here how the difference in abundance of OTU1 or the clustering was generated. Following the argument above that microbes are taken up from the environment, we can think of two scenarios that could generate this pattern. In the first scenario, the plum substrate although heterogeneous, falls into two different categories from a microbial composition perspective. These categories could be states of decay or states that result from the dynamic interplay of microbial metabolites (21). In the second scenario, we might have a cohort of flies that entered the population only recently and brought microbes with them from the previous substrate. We can not disentangle these options at the moment. Similarly, because D. melanogaster communities change with age (18, 22, 23), distinct age cohorts in the population could cause distinct microbial communities. Finally, because flies shape their associated microbial communities (24) and fly host genetic makeup affect the community composition at least under laboratory conditions (4, 5, 13) genetic differences in the host population could play a role in generating the observed pattern.

Because the sample of 5 flies from the plum population clustered with the smaller group consisting of 10 individual flies, it did not represent beta diversity of the whole population well.

## Conclusion

While average bacterial alpha diversity in a *D. melanogaster* population was well represented by a sample of 5 flies, composition of microbial communities is highly variable between individuals from the same population. Larger samples of individual flies could be used to better represent beta diversity in a population. The two clusters we found in our sample of individuals collected from the same substrate, at the same time and location suggests that there are important factors that shape natural *D. melanogaster* microbial communities that we do not understand yet. Because understanding natural communities is what matters for understanding *D. melanogaster* evolution, more research is needed to better understand the factors that shape natural *D. melanogaster* associated microbial communities.

## Material and Methods

### Fly samples

Flies were collected as described previously (12). In short, live flies were collected and brought to the lab in empty vials within 5 hours of collection. Male *D. melanogaster* were identified based on morphology and frozen at -80°C until DNA extraction. Samples from oranges, peaches, and apple1 were the same as in (12). See Table S1 for a full list of sampling locations.

### DNA Extraction, PCR and sequencing

DNA was extracted either from individual male *D. melanogaster* or pools of five males, with the exception *D. melanogaster* orange sample 3 (orange3), for which we were able to retrieve three males only. DNA extraction was performed using the Qiagen QIAamp DNA extraction kit (Qiagen, Carlsbad, CA) using bead beating as described in (12) and running negative extraction controls (without fly material) in parallel.

Barcoded broad range primers, 515F and 806R, as described in Caporaso et al. 2010 (25) were used to amplify the V4 region of the bacterial 16S rRNA gene. DNA was amplified using Phusion^®^ Hot Start DNA Polymerase (Finnzymes, Espoo, Finland) and the following cycling conditions: 30 sec at 98°C; 30 cycles of 9 sec at 98°C, 60 sec at 50°C, and 90 sec at 72°C; final extension for 10 min at 72°C. In order to reduce PCR bias, amplification reactions were performed in duplicate and pooled. PCR products were run on an agarose gel for quantification and pooled in equimolar amounts. Extraction control PCRs were negative and excluded. The resulting pool was gel extracted using the Qiaquick gel extraction kit (Qiagen, Carlsbad, CA) and sequenced on an illumina MiSeq sequencer reading 2 x 250bp.

### Data analysis

Sequencing data was analyzed using mothur (26)(v 1.36.0) following the MiSeq SOP on mothur.org. Sequences were taxonomically classified using the SILVA reference database (27) as implemented in mothur. The ecodist (28) R package was used to calculate Bray-Curtis-Distances. The vegan R package (29) was used for Jaccard and Unifrac distances incorporating the GUniFrac package (30). The pvclust package was used for cluster analysis (31). A detailed analysis script with all mothur and R commands can be found in file S5.

## Data availability

Raw data is available at ncbi SRA under the accession number XX

## Acknowledgements

This work was supported by DFG Forschungsstipendium 1154/1-1 and DFG grant 1154/2-1. We thank Dmitri A Petrov (Stanford University) in whose lab data was collected. We thank Alan O. Bergland and Heather Machado for support in fly sampling. We thank Till Bayer for helpful comments on the manuscript. The authors declare no conflict of interest.

---

**Figure S1.**
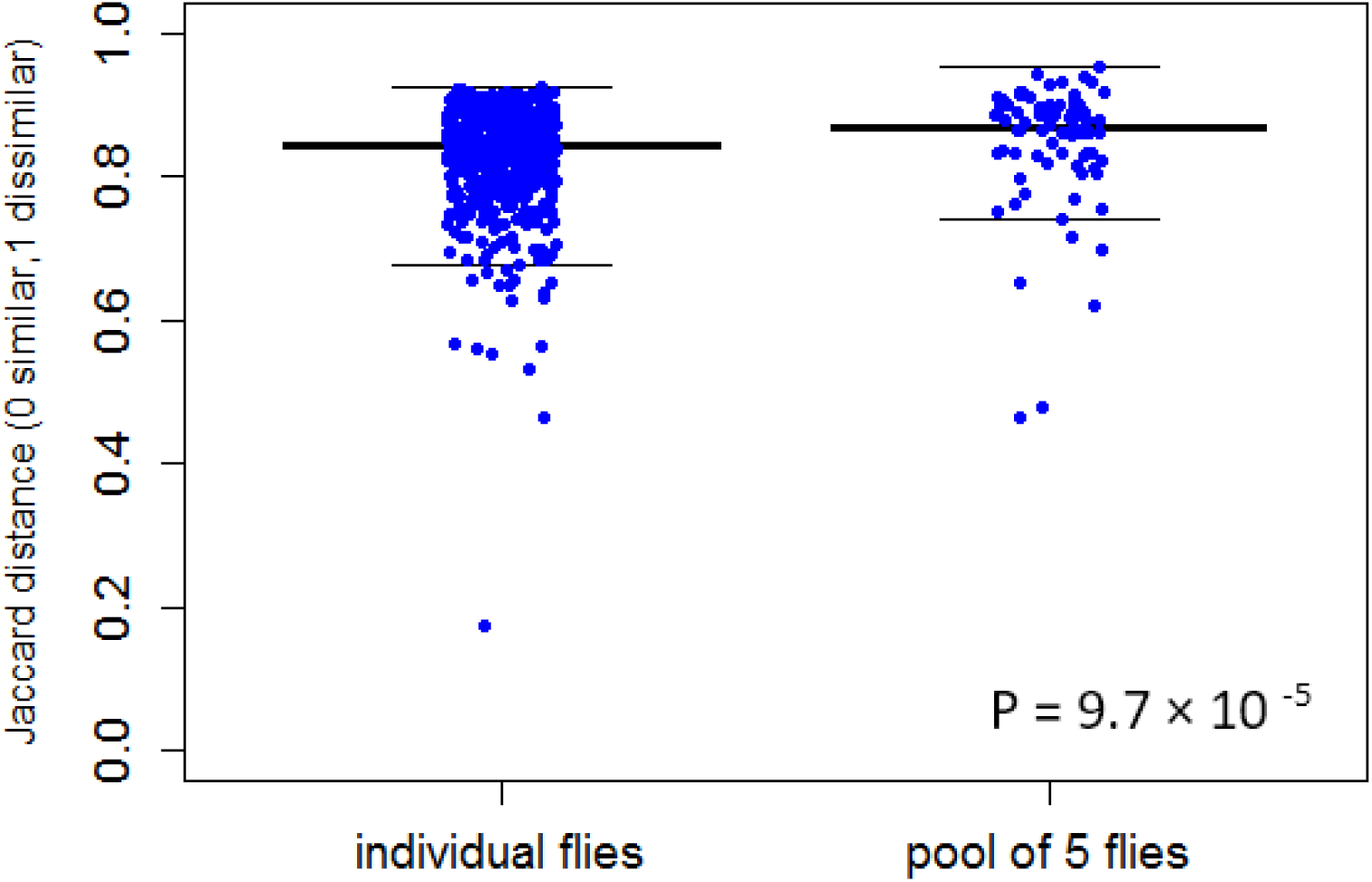
Pairwise Jaccard distances for individual and pooled samples . P-value was computed using Mann-Whitney test.

**Figure S2.**
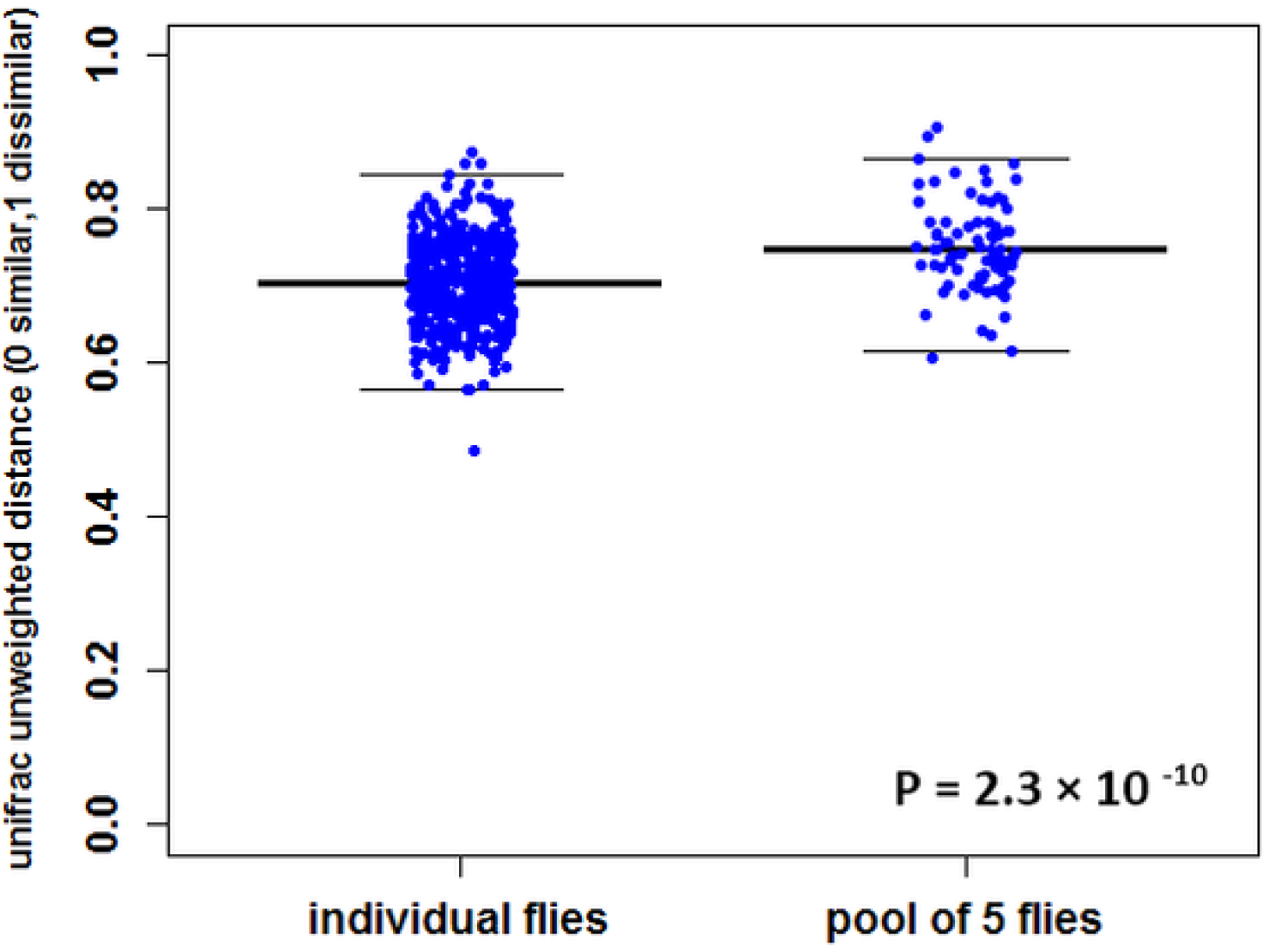
Pairwise unweighted Unifrac distances for individual and pooled samples . P-value was computed using Mann-Whitney test.

**Figure S3.**
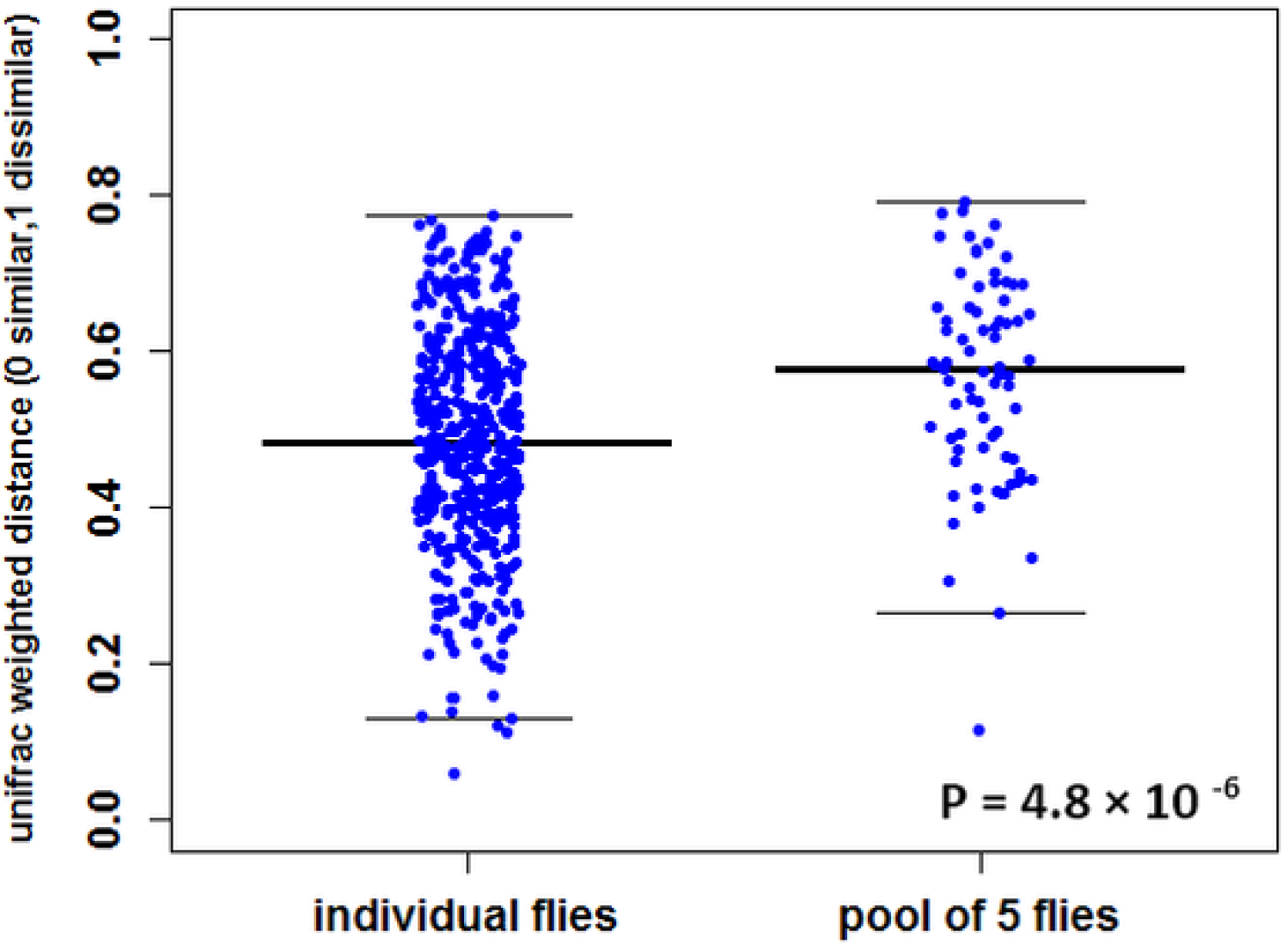
Pairwise weighted Unifrac distances for individual and pooled samples. P-value was computed using Mann-Whitney test.

**Figure S4.**
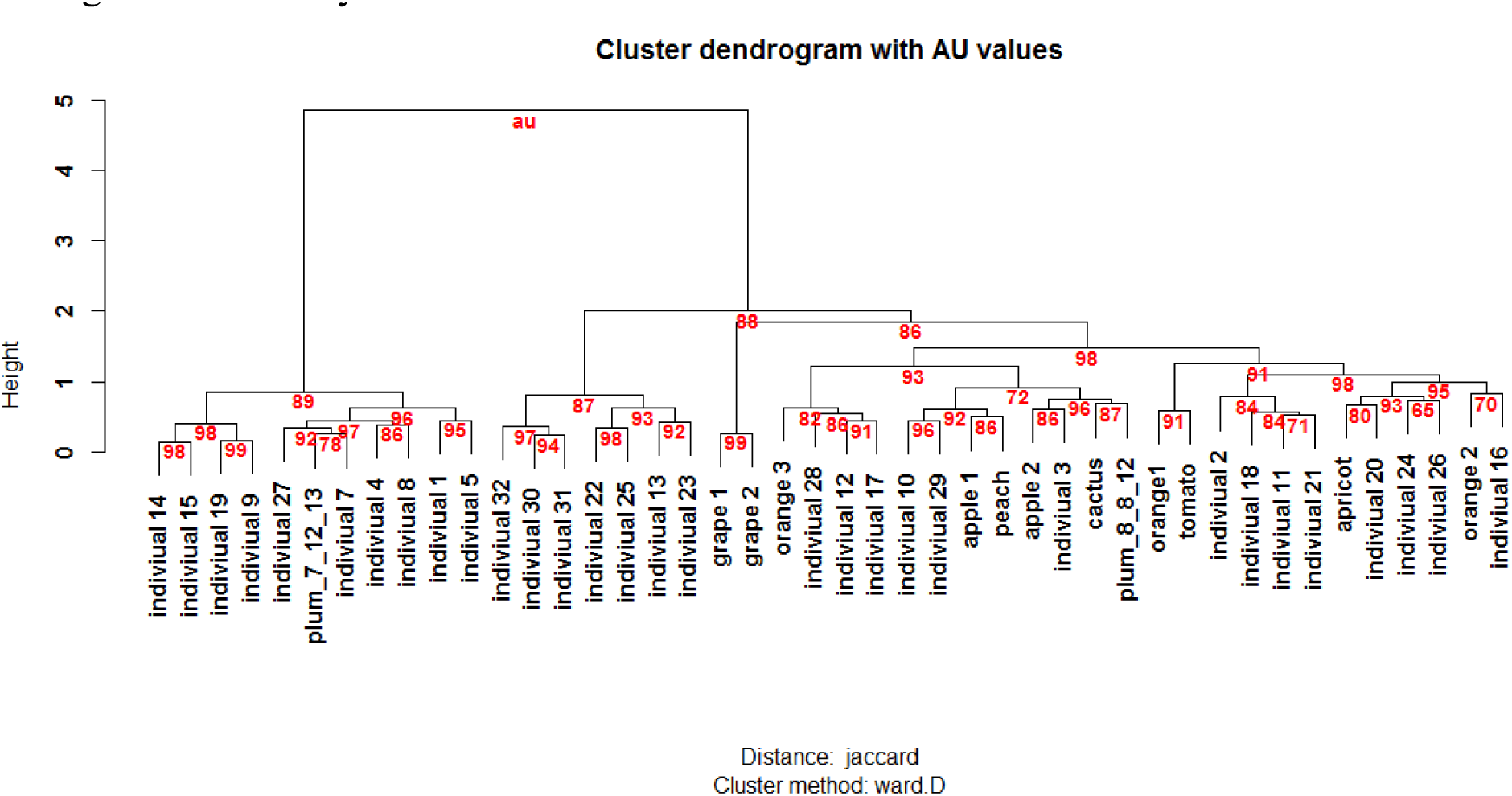
Hierarchical clustering of all samples based on Jaccard distances. Values at branches are AU (Approximately Unbiased) bootstrap support

**Figure S5.**
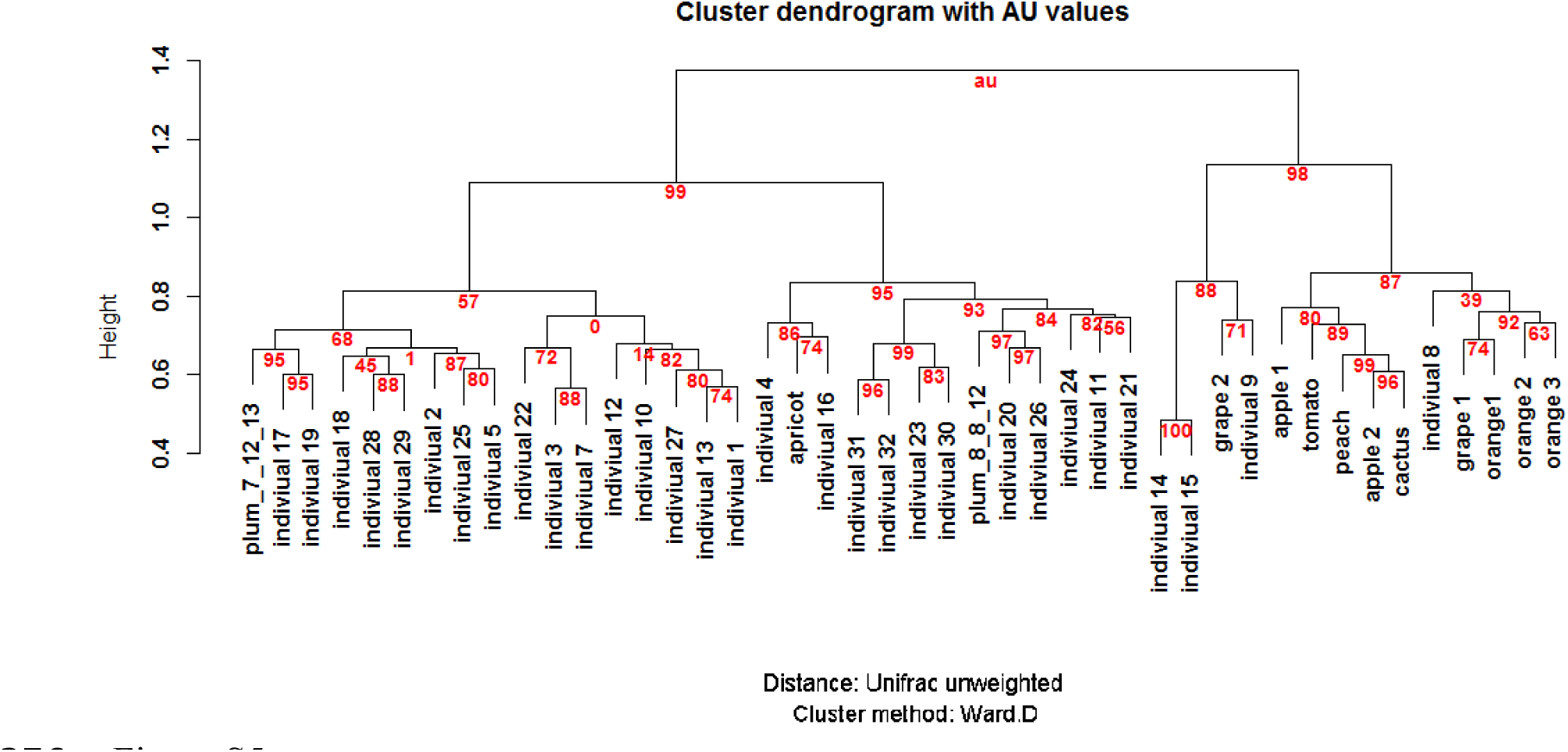
Hierarchical clustering of all samples based on Unifrac unweighted distances. Values at branches are AU (Approximately Unbiased) bootstrap support

**Figure S6.**
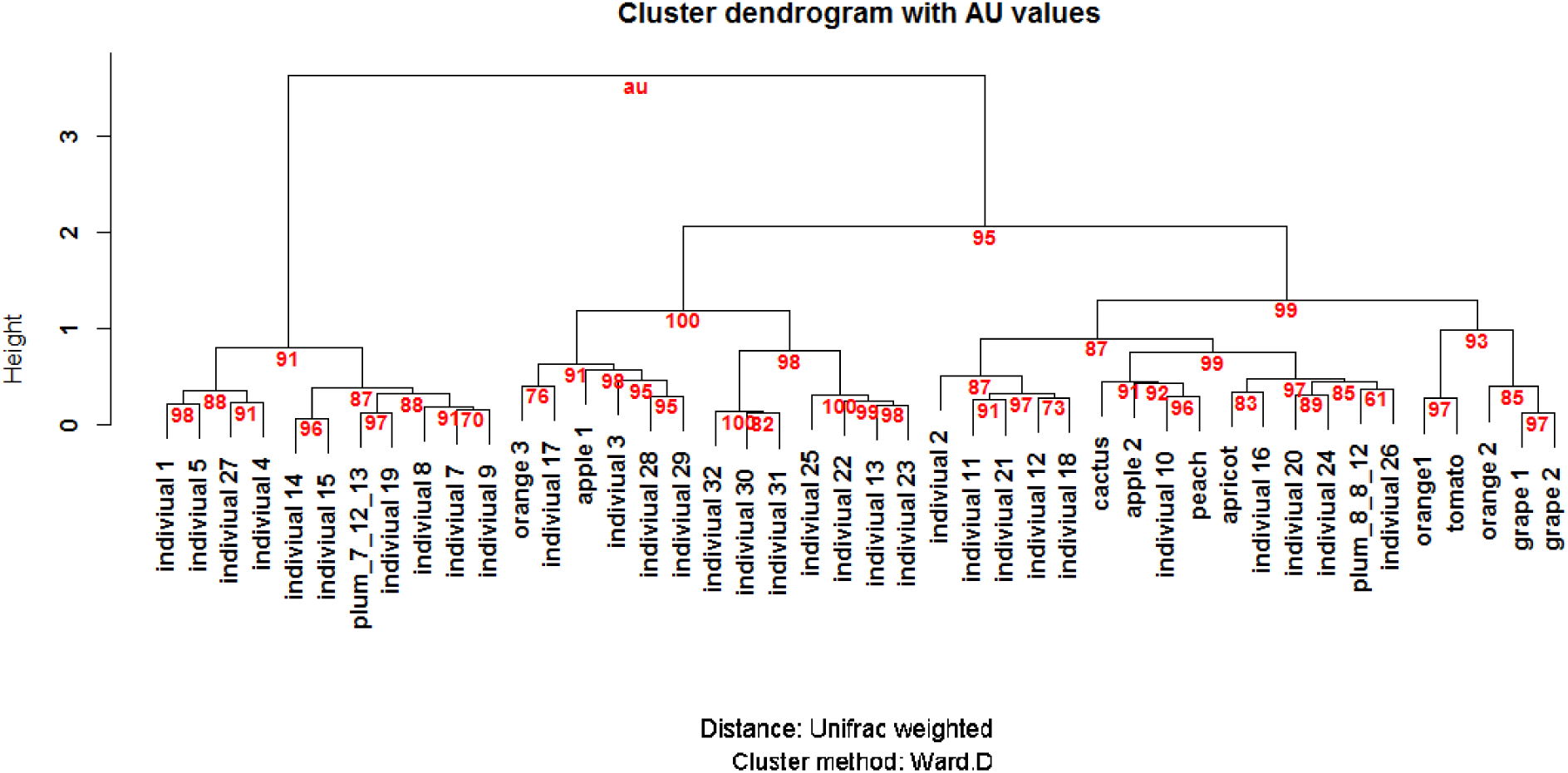
Hierarchical clustering of all samples based on Unifrac weighted distances. Values at branches are AU (Approximately Unbiased) bootstrap support

**Table S1.** list of all samples in the study

| sample | date | location | coordinates |
| --- | --- | --- | --- |
| individual1_7_12_13 | July 12 2013 | Redwood City, CA | 37.48256, -122.23427 |
| individual2_7_12_13 | July 12 2013 | Redwood City, CA | 37.48256, -122.23427 |
| individual3_7_12_13 | July 12 2013 | Redwood City, CA | 37.48256, -122.23427 |
| individual4_7_12_13 | July 12 2013 | Redwood City, CA | 37.48256, -122.23427 |
| individual5_7_12_13 | July 12 2013 | Redwood City, CA | 37.48256, -122.23427 |
| individual6_7_12_13 | July 12 2013 | Redwood City, CA | 37.48256, -122.23427 |
| individual7_7_12_13 | July 12 2013 | Redwood City, CA | 37.48256, -122.23427 |
| individual8_7_12_13 | July 12 2013 | Redwood City, CA | 37.48256, -122.23427 |
| individual9_7_12_13 | July 12 2013 | Redwood City, CA | 37.48256, -122.23427 |
| individual10_7_12_13 | July 12 2013 | Redwood City, CA | 37.48256, -122.23427 |
| individual11_7_12_13 | July 12 2013 | Redwood City, CA | 37.48256, -122.23427 |
| individual12_7_12_13 | July 12 2013 | Redwood City, CA | 37.48256, -122.23427 |
| individual13_7_12_13 | July 12 2013 | Redwood City, CA | 37.48256, -122.23427 |
| individual14_7_12_13 | July 12 2013 | Redwood City, CA | 37.48256, -122.23427 |
| individual15_7_12_13 | July 12 2013 | Redwood City, CA | 37.48256, -122.23427 |
| individual16_7_12_13 | July 12 2013 | Redwood City, CA | 37.48256, -122.23427 |
| individual17_7_12_13 | July 12 2013 | Redwood City, CA | 37.48256, -122.23427 |
| individual18_7_12_13 | July 12 2013 | Redwood City, CA | 37.48256, -122.23427 |
| individual19_7_12_13 | July 12 2013 | Redwood City, CA | 37.48256, -122.23427 |
| individual20_7_12_13 | July 12 2013 | Redwood City, CA | 37.48256, -122.23427 |
| individual21_7_12_13 | July 12 2013 | Redwood City, CA | 37.48256, -122.23427 |
| individual22_7_12_13 | July 12 2013 | Redwood City, CA | 37.48256, -122.23427 |
| individual23_7_12_13 | July 12 2013 | Redwood City, CA | 37.48256, -122.23427 |
| individual24_7_12_13 | July 12 2013 | Redwood City, CA | 37.48256, -122.23427 |
| individual25_7_12_13 | July 12 2013 | Redwood City, CA | 37.48256, -122.23427 |
| individual26_7_12_13 | July 12 2013 | Redwood City, CA | 37.48256, -122.23427 |
| individual27_7_12_13 | July 12 2013 | Redwood City, CA | 37.48256, -122.23427 |
| individual28_7_12_13 | July 12 2013 | Redwood City, CA | 37.48256, -122.23427 |
| individual29_7_12_13 | July 12 2013 | Redwood City, CA | 37.48256, -122.23427 |
| individual30_7_12_13 | July 12 2013 | Redwood City, CA | 37.48256, -122.23427 |
| individual31_7_12_13 | July 12 2013 | Redwood City, CA | 37.48256, -122.23427 |
| individual32_7_12_13 | July 12 2013 | Redwood City, CA | 37.48256, -122.23427 |
| poolof5_plum_8_8_12 | Aug 8 2012 | Redwood City, CA | 37.48256, -122.23427 |
| poolof5_plum_7_12_13 | July 12 2013 | Redwood City, CA | 37.48256, -122.23427 |
| apple2 | Nov 17 2012 | Camino, CA | 38.762123, -120.715358 |
| apricot | July 27 2011 | Guinda, CA | 38.859975, -122.209494 |
| grape1 | Nov 15 2012 | Acampo, CA | 38.189685, -121.231378 |
| cactus | Nov 15 2013 | Stockton, CA | 38.020093, -121.203840 |
| grape2 | Nov 15 2012 | Lodi, CA | 38.138774, -121.209475 |
| tomatoes | July 27 2011 | Capay, CA | 38.708438, -122.075828 |
| orange1 | March 2010 | Manteca, CA | 37.79766, -121.13014 |
| orange2 | Feb 2010 | Escalon, CA | 37.79702, -120.99369 |
| orange3 | March 2010 | Knightsen, CA | 37.97451, -121.65161 |
| apple1 | August 2010 | Johnston, RI | 41.81636, -71.47705 |
| peach | August 2010 | Johnston, RI | 41.81636, -71.47705 |

